# The effect of pre-analytical and physiological variables on cell-free DNA fragmentation

**DOI:** 10.1101/2021.09.17.460828

**Authors:** Ymke van der Pol, Norbert Moldovan, Sandra Verkuijlen, Jip Ramaker, Dries Boers, Wendy Onstenk, Johan de Rooij, Idris Bahce, D. Michiel Pegtel, Florent Mouliere

## Abstract

Assays that account for the biological properties and fragmentation of cell-free DNA (cfDNA) can improve the performance of liquid biopsy. However, pre-analytic and physiological differences between individuals on fragmentomic analysis are poorly defined.

We analyzed the impact of collection tube, plasma processing time and physiology on the size distribution of cfDNA, their genome-wide representation and sequence diversity at the cfDNA fragment-ends using shallow Whole Genome Sequencing.

We observed that using different stabilizing collection tubes, or processing times does not affect the cfDNA fragment sizes, but can impact the genome-wide fragmentation patterns and fragment-end sequences of cfDNA. In addition, beyond differences depending on the gender, the physiological conditions tested between 63 individuals (age, body mass index, use of medication and chronic conditions) minimally influenced the outcome of fragmentomic methods.

Our results highlight that fragmentomic approaches have potential for implementation in the clinic, pending clear traceability of analytical and physiological factors.

## Introduction

Mutation analysis of circulating tumor DNA (ctDNA) is on the path to clinical implementation for guiding treatment choice in oncology [1]. The portfolio of potential applications is increasing and directed towards assessing treatment and pathological response. The impact of pre-analytical variables on the ctDNA tumor fraction and mutation analysis has been intensely investigated [2–6]. The choice of tubes with preservative additives for stabilizing plasma ctDNA in multi-centric studies is now well accepted [1]. The influence of other pre-analytical variables on ctDNA tumor fraction, for example the centrifugation protocol during plasma or urine preparation, remain to be more intensively explored [7].

New applications have recently emerged leveraging the biological and structural properties of cell-free DNA (cfDNA) [8–11]. In plasma samples from healthy individuals, cfDNA is mostly distributed in sizes around a mode of 167 bp, and multiple thereof, suggesting an apoptotic origin [12, 13]. In cancer, and other pathologies, this fragment size distribution can be altered [14–16]. In addition, the cleavage of a given stretch of DNA during cell death is influenced by the chromatin compaction, resulting in non-random fragmentation of cfDNA [11, 17, 18]. Nucleases with characteristic substrate sequences can cleave DNA with distinct nucleosome organization [19]. DNase-1, with a predominant T cleavage site is hypothesized to prefer regions unprotected by nucleosomes, while other enzymes, such as DNase-1L3, with a C cleavage site can act on more tightly packed DNA [20]. Thus, the cleavage sites could be leveraged to map chromatin densities, improve the detection of cancer and to gain information on the tissue localization of the tumor [21]. Previous reports characterize the impact of DNA processing on cfDNA size using PCR methods [22], or electrophoresis measurement [6], but does not assess their influence on other fragmentomic features (cfDNA fragment size distribution, genome-wide cfDNA fragmentation patterns and cfDNA fragment-ends). Despite this rising interest in the field of fragmentomics, the impact of physiological variables on cfDNA fragmentomic features remain mostly unknown.

Here, we investigate the impact of tube collection, plasma processing and preparation on a range of cfDNA fragmentomic analysis (cfDNA fragment sizes, genome-wide patterns and fragment-ends) using low coverage Whole Genome Sequencing. We evaluate with the same methods the impact of physiological variables on cfDNA fragmentomic parameters.

## Results

### 1. Study design

In total, 83 samples from 63 individuals were collected for this study (**Figure 1**). Due to the presence of comorbidities altering cfDNA fragmentation, 1 individual with basal cell carcinoma from the healthy cohort was excluded (**Supplementary table 1** and **Supplementary table 2**). Another 4 healthy individuals were excluded due to missing information related to their physiological conditions (**Supplementary table 2**). 78 samples from 58 individuals were included for analysis. This included 6 lung cancer patients with advanced stage disease. Blood samples were acquired prior to any treatment (**Figure 1** and **Supplementary table 1**). In addition, 52 healthy controls from 2 collection sites were included. Unless otherwise indicated for a specific testing (e.g. different collection tubes), all samples have been processed using the same protocol and conditions (see Methods and **Figure 1**). To compare the collection tubes, 6 lung cancer samples and 20 healthy control samples were used (due to the availability of paired EDTA and Norgen/PAXgene tubes). To assess the impact of processing delay, 30 healthy controls with available recording of the collection, processing and storage time were used. Physiological records including age, gender, body mass index (BMI), comorbidity, use of medication, intoxication (alcohol, drugs), and fasting were available for 42 healthy individuals (**Supplementary table 2**).

**Figure 1:**
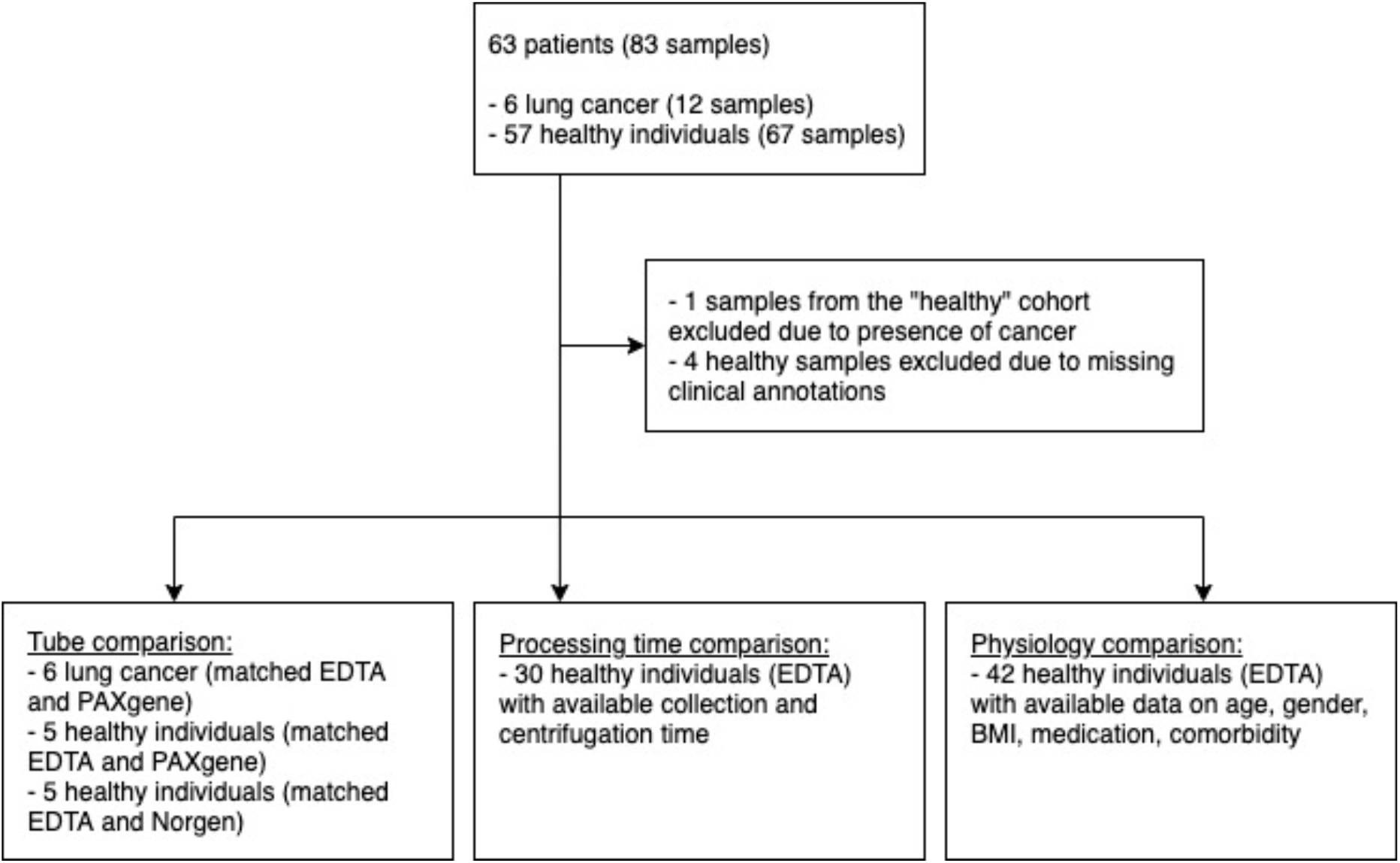
flowchart of the samples collected depending on the parameters tested.

### 2. cfDNA fragmentation differs depending on the blood collection tube

In order to evaluate the impact of the collection tube on cfDNA fragmentation and fragment-end sequence, we selected 16 individuals (10 healthy controls and 6 patients with lung cancer) from whom blood samples were collected both in K2 EDTA pre-coated tubes and in tubes designed for transportation of blood and stabilization of intracellular content (from 2 commercial entities) (**Supplementary Table 1**). In total we processed samples from 32 blood tubes. cfDNA was then isolated using magnetic bead. The concentration in cfDNA was higher in EDTA collected samples vs Norgen or PAXgene collected samples in both healthy and cancer cases, but without a significantly increase (respectively, p-value = 1 and p-value = 0.12, paired Wilcoxon rank sum test) (**Supplementary Figure 1A**). Sequencing libraries were prepared and sequenced at low depth with a low coverage WGS approach (<5-fold coverage). The tumor fraction in ctDNA was calculated from the copy number aberration profiles generated using the ichorCNA software (**Supplementary Figure 1B**). We analyzed paired-end shallow WGS (sWGS) sequencing data to retrieve the cfDNA fragment size distribution (**Figure 2A** and **Supplementary Figure 2**). In all cases we observed the typical cfDNA fragment size distribution centered around a mode of 167 bp. No differences in the cfDNA fragmentation profiles were apparent depending on collection tube (**Supplementary Figure 4**). We calculated the proportions of fragments between 100 and 150 bp (P100_150 bp) and observed no significant differences between EDTA and PAXgene collected samples from healthy individuals (p-value = 0.19, paired Wilcoxon-test), between EDTA and PAXgene collected samples from lung cancer patients (p-value = 0.062, paired Wilcoxon-test), and between EDTA and Norgen collected samples from healthy individuals (p-value = 0.094, paired Wilcoxon-test) (**Figure 2B**), (**Supplementary Table 1**). A more in-depth analysis of the fragmentation per bin size of 30 bp indicate that the differences could be pronounced on specific size ranges depending on the group evaluated (**Supplementary Figure 3**).

**Figure 2:**
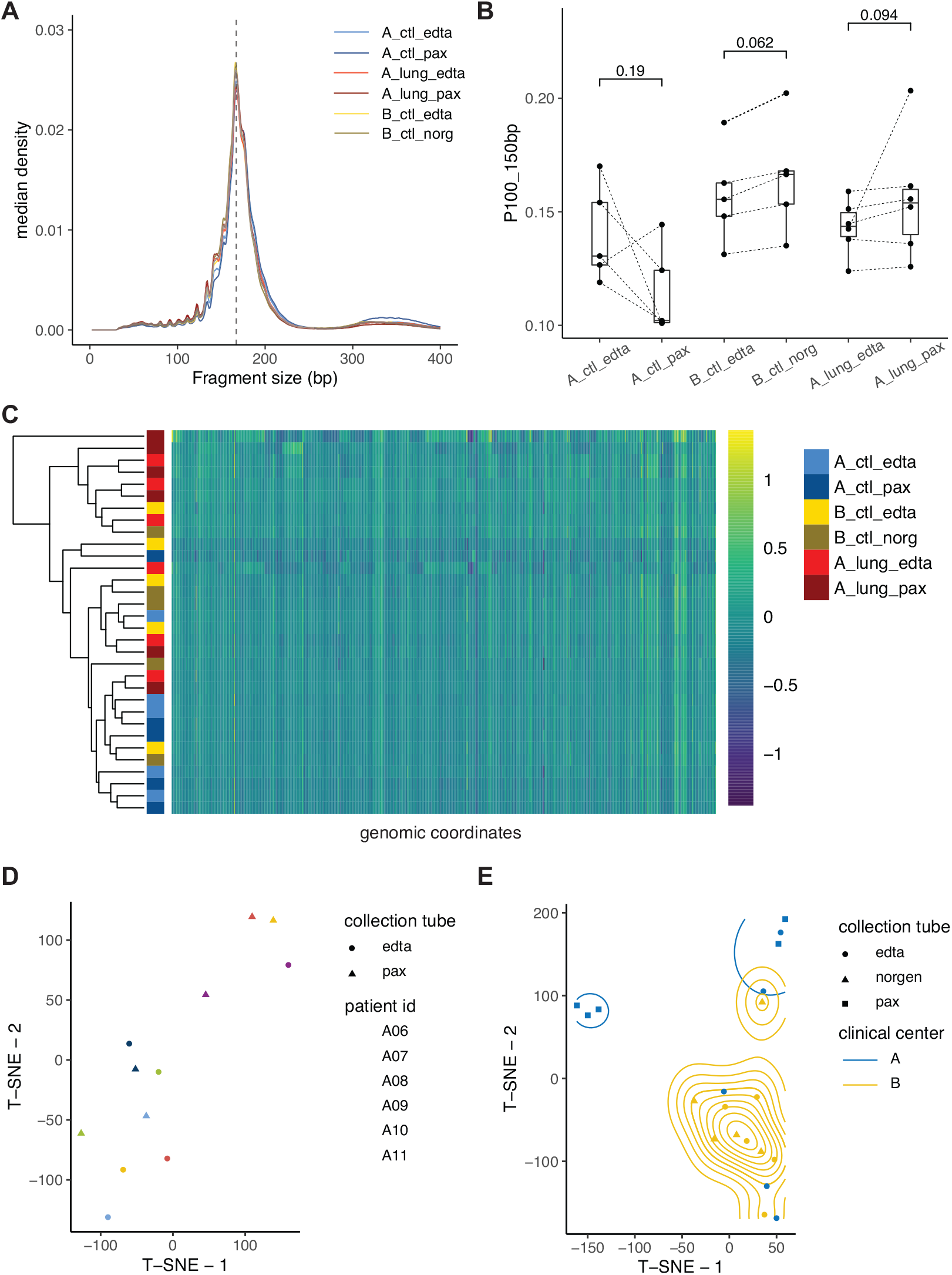
cfDNA fragment sizes and genome-wide pattern depending on the collection center and type of blood tube used. The origin of samples is indicated (clinical center A and B), as well as the type of sample (ctl = healthy individual, lung = lung cancer) and the type of tube used to collect blood (edta = EDTA, norg = Norgen, pax = PAXgene). **A**: Comparison of the median plasma cfDNA size distribution determined by paired-end sWGS. **B:** Proportion of fragments between 100 and 150 bp (P100_150 bp) retrieved from the sWGS data. P-values of the paired Wilcoxon-test comparing the boxplots are added. **C:** Heatmap representing the genome-wide fragmentation profiles of cfDNA samples. In each 5 Mb bin, the z-score log_2_(P100_150 bp / P151_200 bp) was calculated. **D:** A t-distributed stochastic neighbor embedding (t-SNE) plot generated using the Genome-wide cfDNA fragmentation on the cancer samples. **E:** A t-SNE plot generated using the genome-wide cfDNA fragmentation patterns for healthy cases from both clinical centers.

After analyzing the impact of the collection tube on cfDNA fragmentation, we evaluated whether the pre-analytical condition could impact more and notably the genome-wide cfDNA fragmentation patterns [23]. We investigated the size distribution of cfDNA fragments in a position-dependent manner along the genome (**Figure 2C**), (**Supplementary Table 3**). We split the genome into 5 Mb bins and calculated, for each bin, the log2 fraction of short to long fragments (short fragments = 100-150bp; long fragments = 151-220bp) resulting in genome-wide fragmentation profiles for each cfDNA sample (see Methods). In addition, the size distribution of cfDNA fragments was calculated per chromosome arm bin. No difference was observed between the types of collection tube on chromosome arm binned fragmentation profiles (**Supplementary Figures 6-8**). Inputted in unsupervised clustering, namely a t-distributed stochastic neighbor embedding (t-SNE), matched EDTA and PAXgene collection tubes of the lung cancer patients tend to cluster together (**Figure 2D**). As for the healthy samples, clustering was based on the sample source rather than type of collection (**Figure 2E**). The use of cfDNA stabilizing tubes (PAXgene, Norgen) does not alter this clustering, suggesting that multi-centric studies using stabilizing tubes could also be affected by potential bias in fragmentomic studies (**Figure 2E**).

Next, we calculated the proportions of cfDNA fragments ending in a specific trinucleotide sequence from the same sequencing data (see Methods) (**Figure 3A**), (**Supplementary table 4**). We observed that the trinucleotide CCA, CCT and CCC are overrepresented in both healthy controls and cancer cases as previously described [20]. The fragment-end diversity (calculated using a normalized Shannon entropy index) was comparable between healthy samples collected in EDTA and either PAXgene or Norgen tubes (respectively, p-value=0.81 and p-value=0.12, paired Wilcoxon-test) (**Figure 3B**). Similarly, the fragment-end diversity for lung cancer samples collected using EDTA or PAXgene tubes is comparable (p-value=0.84, paired Wilcoxon-test) (**Figure 3B**). Using a t-SNE on the cfDNA fragment-end trinucleotide fractions, we observed that matched EDTA and PAXgene collection tubes originating from the same lung cancer patient clustered together (**Figure 3C**). The impact of sample source identified using genome-wide cfDNA patterns was confirmed by unsupervised clustering of the cfDNA fragment-end trinucleotide fractions. Despite undergoing the same processing procedure, healthy control samples originating from clinical center B clustered together (either if they were collected in EDTA tubes or Norgen tubes) but differently from the healthy controls of clinical center A (collected in EDTA tubes or PAXgene tubes) (**Figure 3D**).

**Figure 3:**
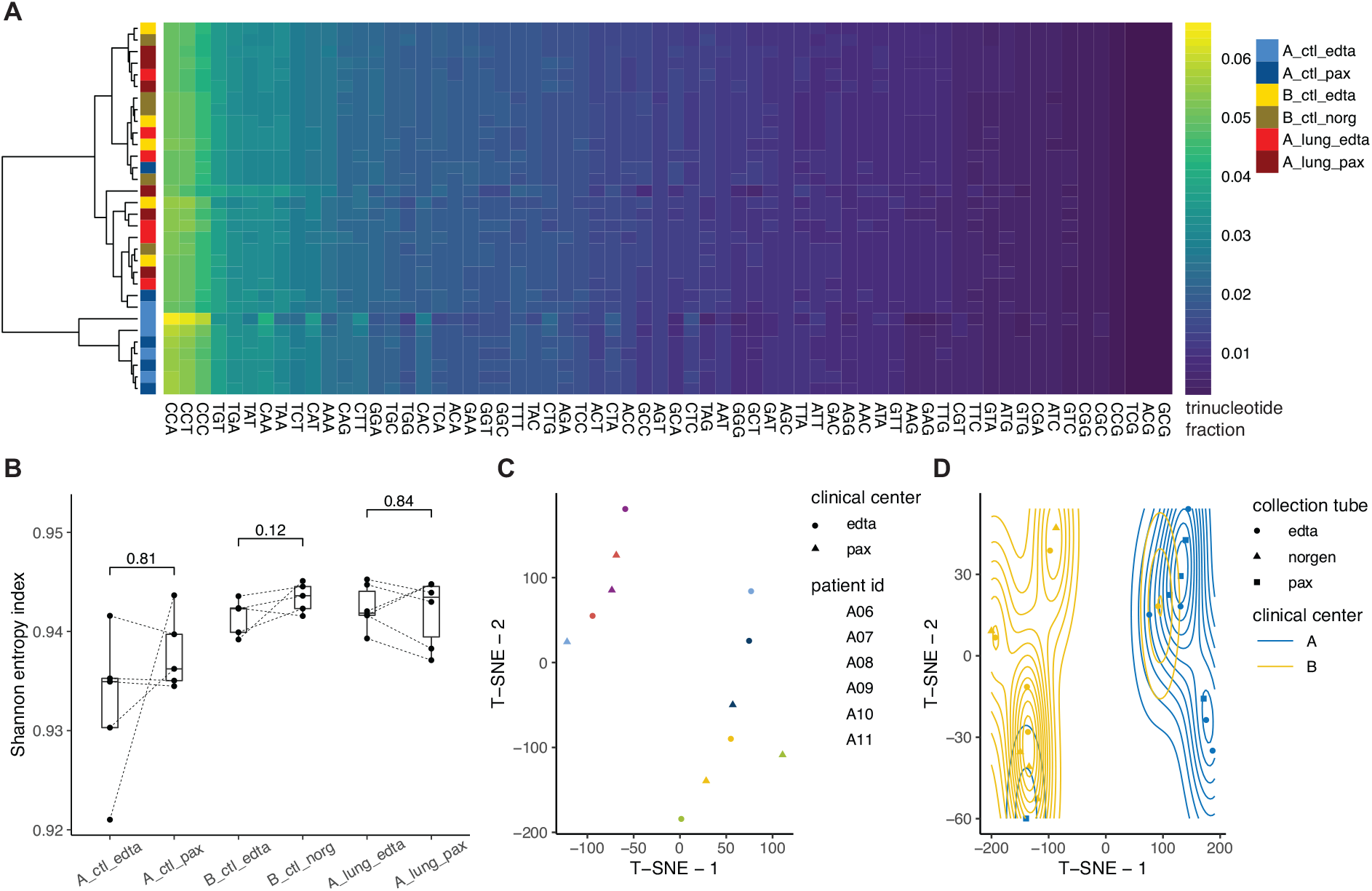
cfDNA fragment-ends do not differ depending on the type of collection tube used but are affected by the center of collection. The origin of samples is indicated (clinical center A and B), as well as the type of sample (ctl = healthy individual, lung = lung cancer) and the type of tube used to collect blood (edta = EDTA, norg = Norgen, pax = PAXgene). **A**: Heatmap of the proportions in trinucleotides at the end of cfDNA fragments recovered by paired-end sWGS analysis. The color intensity represents the proportion in fragments (low in blue, high in yellow). The annotation on the left side indicates the source of the samples, their type and tube used. Rows are clustered using a Ward algorithm. **B**: Diversity in fragment-ends of cfDNA are calculated using a Shannon entropy index depending on the origin of the samples, the type of sample and the type of tube. P-values of the paired Wilcoxon-test comparing the boxplots are added. **C**: A t-SNE plot, generated using the cfDNA fragment-end trinucleotides for cancer samples. **D:** A t-SNE plot generated using the cfDNA fragment-end trinucleotides for healthy samples with associated density contours.

### 3. Impact of processing time on cfDNA fragmentation features

The transportation of blood samples and delayed processing times have been reported to be a confounding factor for mutation analysis using cfDNA [2, 3]. To analyze the impact of the processing time (from the blood collection, to the isolation of plasma and storage in a freezer), we recorded the processing time for a cohort of 30 healthy individuals using EDTA tubes (**Figure 1** and **Supplementary table 2**). Samples from 12 additional individuals where processing time was not recorded have been added to the comparison. Samples were prepared and sequenced using the same sWGS approach as earlier described (see Methods), and multiple fragmentomic features were recovered from the sequencing data. We do not observe a significant difference of the processing time on cfDNA fragmentation pattern using either the P100_150 bp (**Figure 4A**) or on the diversity of fragment-ends using the Shannon entropy index (**Figure 4B**). Genome-wide cfDNA fragmentation showed minor differences between processing time groups depending on chromosome arm (**Supplementary Figure 8**).

**Figure 4:**
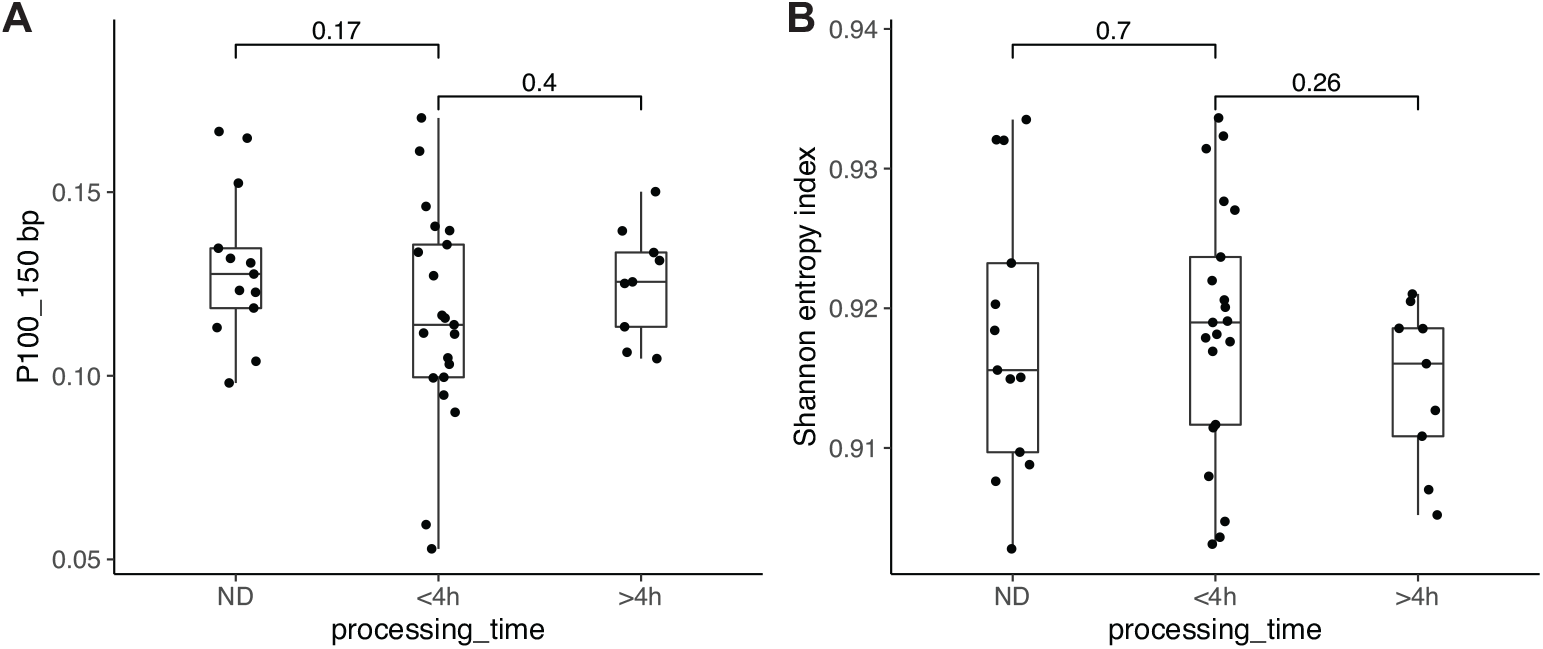
cfDNA fragmentation and fragment-end diversity does not differ depending on the plasma processing time. The processing time is expressed in hours and corresponds to the time required to process the samples from the blood collection to the storage of the plasma in a freezer. N.D.: processing time not determined or recorded. P-values of the Wilcoxon-test comparing the boxplots are added. **A**: Proportion of short cfDNA fragment comprised between 100 and 150 bp (P100_150) depending on the duration of the plasma processing time. **B**: Diversity in fragment-ends of cfDNA calculated using a Shannon entropy index depending on the duration of the plasma processing time.

### 4. Impact of physiological variables on cfDNA fragmentation and fragment-ending in non-cancer controls

In addition to pre-analytical or other technical variables, physiology could alter the fragmentation patterns and other fragmentomic properties of cfDNA. In order to assess the impact of major physiological variables we recruited a cohort of 47 individuals without cancer using EDTA tubes, with an extensive annotation of their conditions. One individual declared pathology, 4 individuals did not provide any clinical information and were excluded from further comparative analysis (**Figure 1** and **Supplementary table 2**). The same protocol to prepare sequencing libraries was applied to all samples (see Methods), and they were sequenced in the same sequencing run. We analyzed paired-end sequencing data to retrieve the cfDNA fragment size distribution (**Supplementary Figure 4**), genome-wide cfDNA fragmentation patterns (**Figure 5A** and **Supplementary Figures 10-14**) as well as the trinucleotide composition of the cfDNA fragment-end sequences. Overall, our cohort of 42 individuals consisted of 13 males and 29 females. The median age of the individuals recruited was 34 years old (range 26-64). The median body mass index was 23.3 (range 16.5-33.9). In our cohort, 8 individuals presented a pathology (or long-term condition) which was reported during the sampling (**Supplementary Table 2**). We also compared whether the cfDNA fragmentomic properties were altered by the usage of medicine (for example contraceptive pills or antihistamins), exercise or alcohol intake (**Supplementary Table 2**).

**Figure 5:**
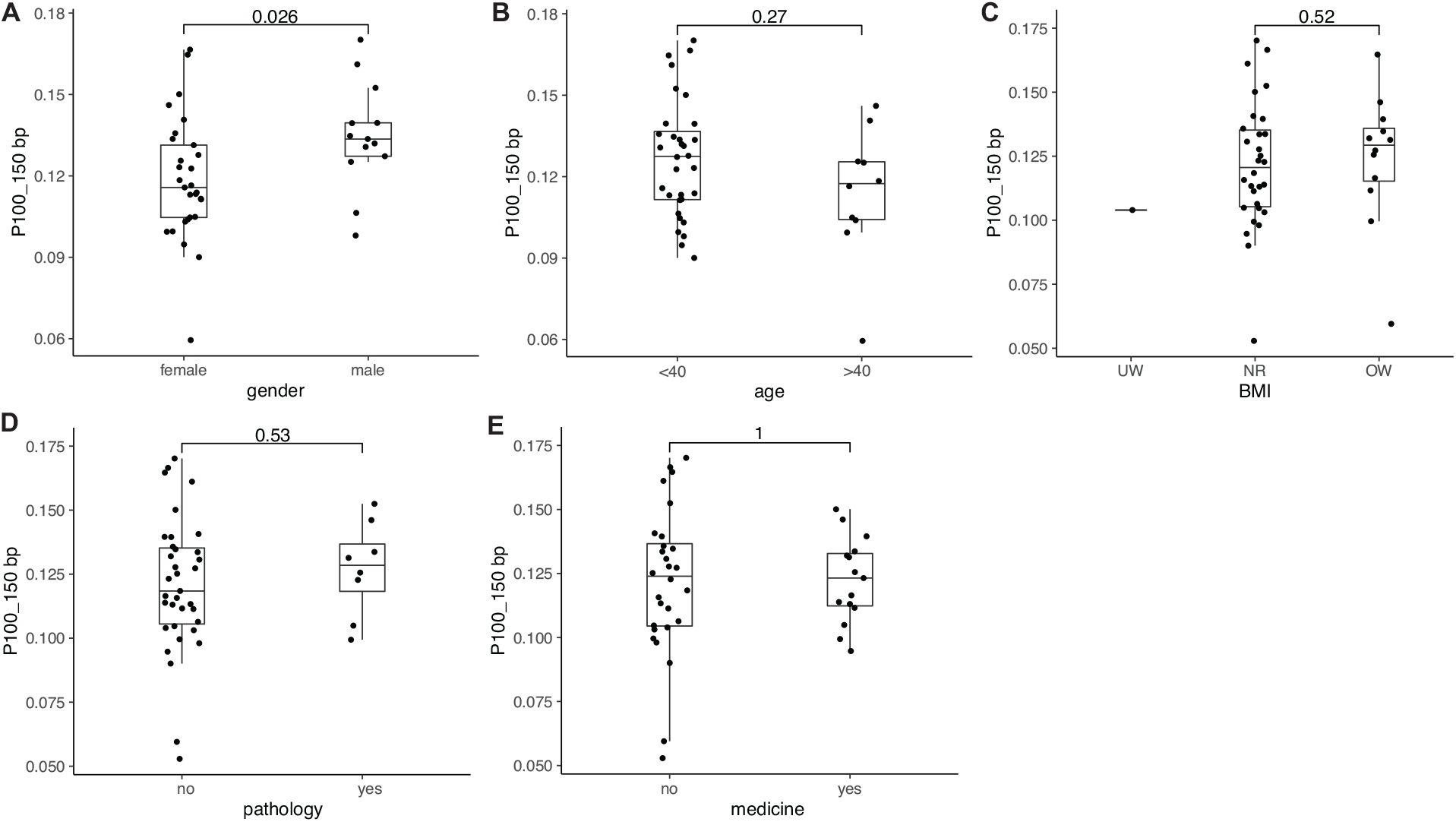
the proportion in short cfDNA differs depending on the gender, but not depending on other physiological parameters. Proportion of short cfDNA fragments (P100_150 bp) for the same samples depending on their gender (**A**), age (**B**), Body Mass Index (BMI) (**C**), presence of non-cancer pathologies (**D**), and medicine taken (**E**). P-values of the Wilcoxon-test comparing the boxplots are added. UW: underweight, NR: Normal weight range, OW: overweight.

The proportion of fragments between 100 and 150 bp is significantly higher in male samples (p-value = 0.026, Wilcoxon-test, **Figure 5A**) (**Supplementary table 2**). The cfDNA fragment size distribution does not significantly differ depending on the age of the individuals in our cohort (p-value = 0.27, Wilcoxon-test, **Figure 5B** and **Supplementary Figure 15**), the BMI (p-value = 0.52 between NR and OW, Wilcoxon-test, **Figure 5C** and **Supplementary Figure 16**), the presence of non-cancer pathologies (p-value = 0.53, Wilcoxon-test, **Figure 5D**) and the use of medication before sampling (p-value = 1, Wilcoxon-test, **Figure 5E**).

Genome-wide cfDNA fragmentation patterns analyzed using 5 Mb bins of the genome does not result in the formation of clusters depending on all tested physiological parameters of the individuals in our cohort (**Figure 6A**) (**Supplementary table 5**). Assessing the genome-wide cfDNA fragmentation patterns on a chromosome arm level revealed minor differences depending on age group, medicine use, the presence of chronic disease and exercise (**Supplementary Figures 10-14**) (**Supplementary table 6**).

**Figure 6:**
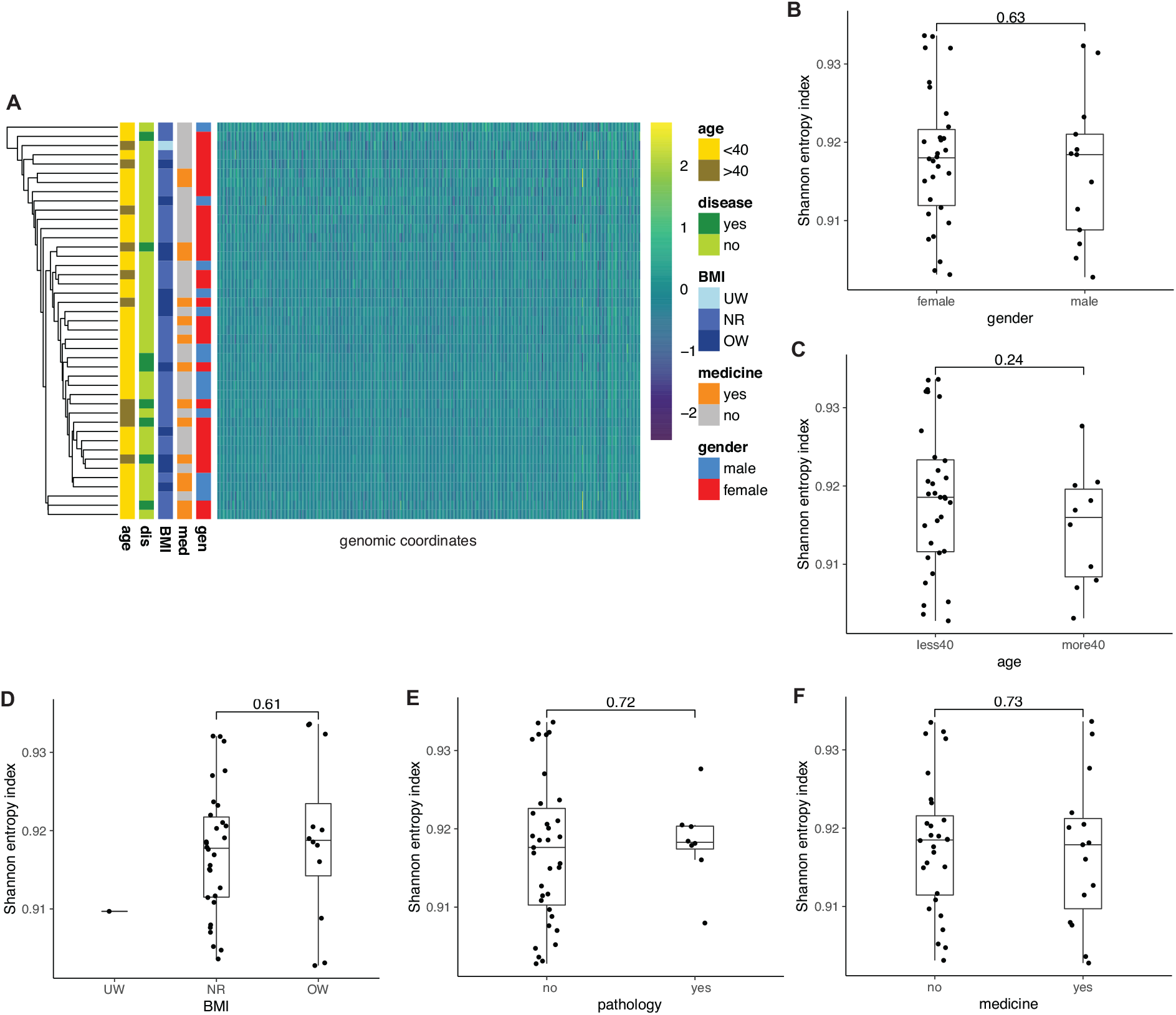
cfDNA genome-wide fragmentation and fragment-ends do not differ depending on the age, gender, body mass index (BMI), presence of non-cancer chronic disease, or medicine taken. Heatmap representing the genome-wide fragmentation profiles of cfDNA samples (**A**). In each 5 Mb bin, the z-score log_2_(P100_150 bp / P151_220 bp) was calculated. Physiological variables are added as columns annotations. Diversity in fragment-end sequences calculated using a normalized Shannon entropy index depending on the gender (**B**), age (**C**), BMI (**D**), presence of non-cancer pathology (**E**), the use of medicine before sampling for the same samples (**F**). P-values of the Wilcoxon-test comparing the boxplots are added.

Regarding the impact of gender on the fragment-end sequences, no significant differences were observed in cfDNA fragment-end diversity, which is calculated using a normalized Shannon entropy index (p-value = 0.63, Wilcoxon-test, **Figure 6B**). The diversity in trinucleotides at the end of cfDNA fragments was not affected by the age of the individuals in our cohort (p-value = 0.24, Wilcoxon-test, **Figure 6C**) (**Supplementary table 7**). The cfDNA fragment-endings were not affected by the presence of non-cancer pathologies (p-value = 0.72, Wilcoxon-test, **Figure 6E**) or the use of medicine before sampling (p-value = 0.73, Wilcoxon-test, **Figure 6F**).

## Discussion

The pre-analytical conditions and physiology of individuals can influence the biological properties and structure of cfDNA, and subsequently the interpretation of these data in clinical studies. Here, with a specific focus on the type of collection tube, the processing time and the physiology of individuals we analyzed the impact of these variables on various fragmentomic approaches: the cfDNA fragment size distribution, genome-wide cfDNA fragmentation patterns and cfDNA fragment-ends.

The choice of the collection tube, and the collection center, impacts certain aspects of fragmentomic analysis. When analyzing the global fragment size distribution of cfDNA, these differences are not clearly observable, confirming that this metric, despite being potentially less sensitive, remains robust irrespective of the collection tube used or the clinical collection center. By focusing on the size differences of cfDNA depending on their genomic coordinates, or by analyzing the composition in bases at the end of cfDNA fragments (as well as their diversity), some differences can be observed. In particular, healthy samples collected from different locations despite being processed with the same preparation protocol, cluster separately. Plasma cfDNA isolated from the stabilizing tubes (PAXgene or Norgen) cluster with their corresponding EDTA samples using both genome-wide cfDNA fragmentation patterns or the composition in bases from the fragment-ends. These results suggest that multi-centric studies using stabilizing tubes to preserve cfDNA could still be affected by batch effect for fragmentomic analysis. This extends to multicentric studies comparing cancer cases against healthy individuals collected across different clinical centers.

Beyond the collection tube, the delay in processing time (from blood collection, to plasma isolation and storage) does not seem to affect significantly the cfDNA fragment size distribution, proportion in short cfDNA fragments (between 100 and 150 bp), or the composition and diversity in bases at the end of cfDNA fragments. We observed a moderate impact of the processing time on the genome-wide cfDNA fragmentation patterns. We caution that we investigated solely the impact of the processing time on samples collected using EDTA collection tubes and not stabilizing tubes (which might reduce the variability observed in some cases). Also, beyond the processing time, we have not investigated the centrifugation speed and protocol used to isolate plasma from whole blood. Previous reports indicated that the centrifugation speed could impact the composition of the plasma in cfDNA and extracellular vesicles [7], and such centrifugation times should be investigated further to properly assess their impact on cfDNA fragmentation.

As it reflects chromatin organization, we could anticipate that cfDNA fragmentation and fragmentomic features could be affected by the physiology of individuals. Previous work indicates that age could modify the observed size of cfDNA fragments using sequencing [24]. We have not observed such a difference in our dataset, even if our conclusion could be hampered by the small proportion of older individuals in our cohort (median age = 34 years old). On a global fragmentation scale, the proportion of short fragments (P100_150 bp) differed depending on gender. This effect was not observed in the analysis of genome-wide cfDNA patterns, which is due to the application of z-score normalization that corrects for global fragmentation artefacts. Fragment-end analysis was not impacted by gender. Previous studies reported a lower cfDNA concentration in females compared to males [25]. Therefore, a potential difference in metabolic activity associated with gender could impact the fraction of short fragments recovered but would not exert an effect on fragment end analysis. The proportion of short fragments recovered was not found to be altered for any other physiological variables. When applying unsupervised clustering to both genome-wide cfDNA fragmentation and fragment-end information, no clusters were formed based on physiological variables.

Our work is limited by the complexity of the fragmentomic parameters and that we restricted our questions to investigate to some clear clinical endpoints. For example, the diversity of fragment-end, their jaggedness as well as the topology of cfDNA are not investigated in this work and would require a specific analysis [26]. In addition, we only evaluate the impact of the pre-analytical and physiological variables on double-stranded DNA (dsDNA). Recent works highlighted the presence of single-stranded or heavily damaged dsDNA, with altered fragmentomic properties, that could be analyzed using only single-stranded sequencing libraries [15, 27]. We also focused our analysis to plasma samples, cfDNA from other bio-fluids might be altered differently by the pre-analytical conditions of collection as well as by the physiology of the patients [28, 29]. The clinical impact of our work is also hampered by the relatively limited number of cases included in this study.

Nevertheless, our results indicate that a comprehensive assessment of the impact of pre-analytical variables and physiological variables is necessary to clearly interpret biological data and improve the clinical implementation of fragmentomic approaches for cfDNA analysis and cancer detection.

## Methods

### Patient and control cohort

Blood samples from healthy individuals have been collected in 2 clinical centers in the Netherlands and processed using the same protocol. Part of the healthy samples (B) were collected in the Urimon Study (NL67854.041.18) and obtained from the Stibion Biobank (Enschede, The Netherlands). They were included in this study by request from You2Yourself BV for quality control purposes. The other blood samples from healthy individuals and lung cancer patients have been collected at the Amsterdam University Medical Center (A) using the same protocol. In total 71 blood samples have been collected from 59 healthy controls, and 12 blood samples from 6 lung cancer patients (**Supplementary table 1** and **Supplementary table 2**). The study has been approved by the ethics board of the Amsterdam UMC (METc 2021.0388). Written informed consent has been obtained from all individuals included in this study.

### Sample collection and DNA isolation

The blood samples from the 2 clinical centers were processed using the same protocol. Whole blood was collected in EDTA K2 collection tubes and plasma isolated within 5.5 hours. Briefly, blood was centrifuged at 900 g for 7 min at room temperature. Supernatant was collected carefully, and centrifuged further at 2500g for 10 min at room temperature. Isolated plasma was aliquoted in 0.5 mL Nunc tubes and stored at -80C. Samples collected in PAXgene tubes were processed following the constructor recommendations. Blood samples were inverted 10 times before processing. Blood was centrifuged at 1900 g for 15 min at room temperature. Supernatant was collected carefully, and centrifuged further at 1900 g for 10 min at room temperature. Isolated plasma was aliquoted in 0.5 mL Nunc tubes and stored at -80C. cfDNA was isolated from all plasma samples using the QIAsymphony Circulating DNA Kit (Qiagen) according to the manufacturer’s instructions. Between 2 and 3.5 mL of plasma was used for the cfDNA extraction. The cfDNA concentration was determined using an Agilent 4200 TapeStation System with the Cell-free DNA ScreenTape Analysis assay (Agilent).

### Library preparation and sequencing

Sequencing libraries were prepared for WGS using the ThruPLEX® Plasma-seq Kit (Takara Bio) according to the manufacturer’s instructions. cfDNA input was 1-10 ng per extracted cfDNA sample. Sequencing libraries quality was controlled using an Agilent 4200 TapeStation System with the D1000 ScreenTape Analysis assay (Agilent). Sequencing libraries were pooled on equimolar amounts and sequenced using 150-bp paired-end runs on the Illumina NovaSeq 6000 using S4 flow-cells (Illumina).

### Whole-genome sequencing data processing

Initial quality control of demultiplexed data was performed using the FastQC software (v0.11.9). Trimming was done using BBDuk (BBmap v38.79). Quality check post trimming was carried out by the FastQC software (v0.11.9). Trimmed reads were aligned to human reference genome GRCh38 with the BWA-MEM software (v0.7.17) using the default settings. MarkDuplicates (Picard Tools v2.22.2) was applied to annotate duplicate reads. Reads with a MAPQ score below 5, PCR duplicates, secondary alignments, supplementary alignments and unmapped reads were removed prior to further downstream analysis using samtools (v1.9). Samtools-flagstat software (v1.9) and qualimap software (v2.2.2) were performed as a post alignment quality check.

### Copy number analysis

The ichorCNA software (commit 5bfc03e) [30] was used to perform the copy number analysis and estimate the ctDNA tumor fraction from bam files. Exceptions to the software’s default settings are as follows: i) An in-house panel of sWGS normals was created, ii) non-tumor fraction parameter restart values were increased to c(0.95,0.99,0.995,0.999), iii) ichorCNA ploidy parameter restart value was set to 2, iv) no states were used for subclonal copy number and v) the maximum copy number to use was lowered to 3. The reported tumor fraction was retrieved from the data using the highest log likelihood solution.

### cfDNA fragment size distribution analysis

To extract information about the cfDNA fragment size distribution, the Picard InsertSizeMetrics software (v2.22.2) was used with the default settings on the mapped reads. For each sample the proportion of fragments between 100 and 150 bp were calculated from sWGS data for comparison purposes. cfDNA fragmentation plots were constructed in R (v3.6) using the packages ggplot (v3.3.5), dplyr (v1.0.7), tidyr (v1.1.3).

### Genome-wide cfDNA fragmentation patterns analysis

The genome-wide cfDNA fragmentation patterns analysis was adapted from a previous report [18] with the following modifications: the GRCh38 reference genome was binned into 5Mb bins. To account for inter-sample short fragment fraction variability, z-score normalization was applied over all 5Mb bins from the same sample. Regional fragmentation plots were constructed in R (v3.6) using the packages ggplot (v3.3.5), pheatmap (v1.0.12), dplyr (v1.0.7), tidyr (v1.1.3) and viridis (v0.6.1).

### Fragment end analysis

Fragment end analysis was performed using the Fragment End Integrated Analysis (FrEIA) tool. In brief, the first 3 mapped bases from the 5’ end of the read pairs was retrieved. The frequency of fragments starting with each of the 64 trinucleotide possibilities was calculated. The normalized Shannon entropy index was computed using the following formula

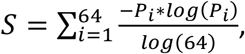

where *P*_*i*_ *is the frequency of a specific trinucleotide ending*. Fragment end plots were constructed in R (v3.6) using the packages ggplot (v3.3.5), pheatmap (v1.0.12), dplyr (v1.0.7), tidyr (v1.1.3) and viridis (v0.6.1).

## Data availability

Raw sequencing data will be deposited upon publication at the European Genome-phenome Archive.

## Author information

## Acknowledgement

The authors are thankful to Mai Tran and the Amsterdam UMC Liquid Biopsy Center for the logistical support and advices. Y.P., W.O. and F.M. are funded by the Amsterdam UMC Liquid Biopsy Center, an initiative made possible through the Stichting Cancer Center Amsterdam. The authors would like to thank Dr. Dineika Chandrananda for comments and discussions to improve the analysis of the ichorCNA algorithm. N.M. and F.M. are supported by a Dutch Cancer Fund (KWF-12822).

## Authors contribution

Conception and design: Y.P., and F.M., Experiments and data collection: Y.P., J.Ramaker, S.V., W.O., Data processing: Y.P., N.M., D.B., F.M., Data analysis: Y.P., N.M., F.M., Sample acquisition: D.M.P., I.B., J.Rooij, Funding acquisition: D.M.P., I.B., F.M., Manuscript draft: Y.P., N.M., F.M., Manuscript revisions and comments: all authors. Supervision: D.M.P. and F.M.

