## Supplementary Figures for "The effect of pre-analytical and physiological variables on cell-free DNA fragmentation"

**Suppl. Figure 1**

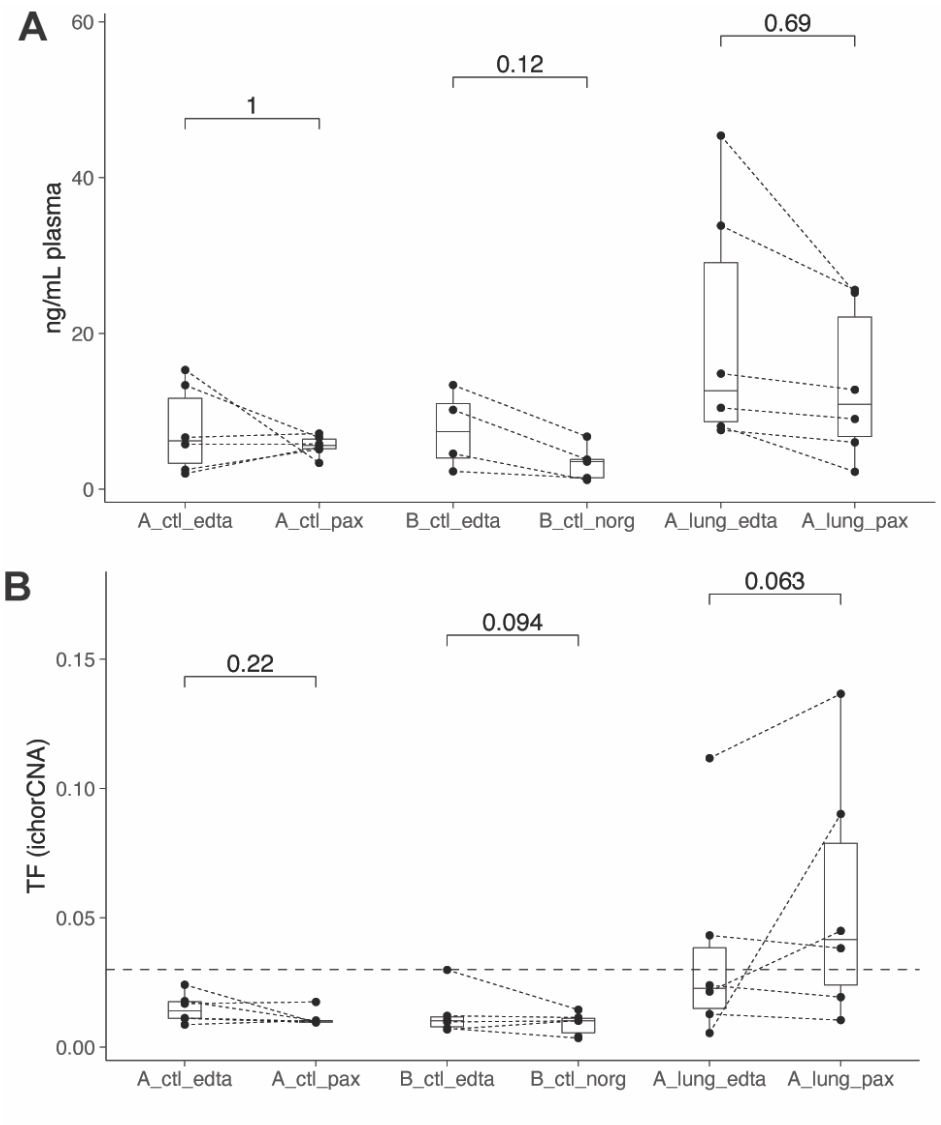


**Suppl. Figure 1: concentration in cfDNA and ctDNA tumor fraction from the matched plasma samples depending on the tube used for collecting blood. A)** Comparison of the concentration in cfDNA (in ng/mL of plasma) depending on the collection tube used on paired samples. The p values are indicated (Wilcoxon paired test). **B)** Comparison of the tumor fraction as estimated by ichorCNA depending on the collection tube (EDTA versus PAXgene or EDTA versus Norgen), collection centre (A vs B) and patient characteristics (cancer versus controls). The p values are indicated (Wilcoxon paired test).

### Suppl. Figure 2

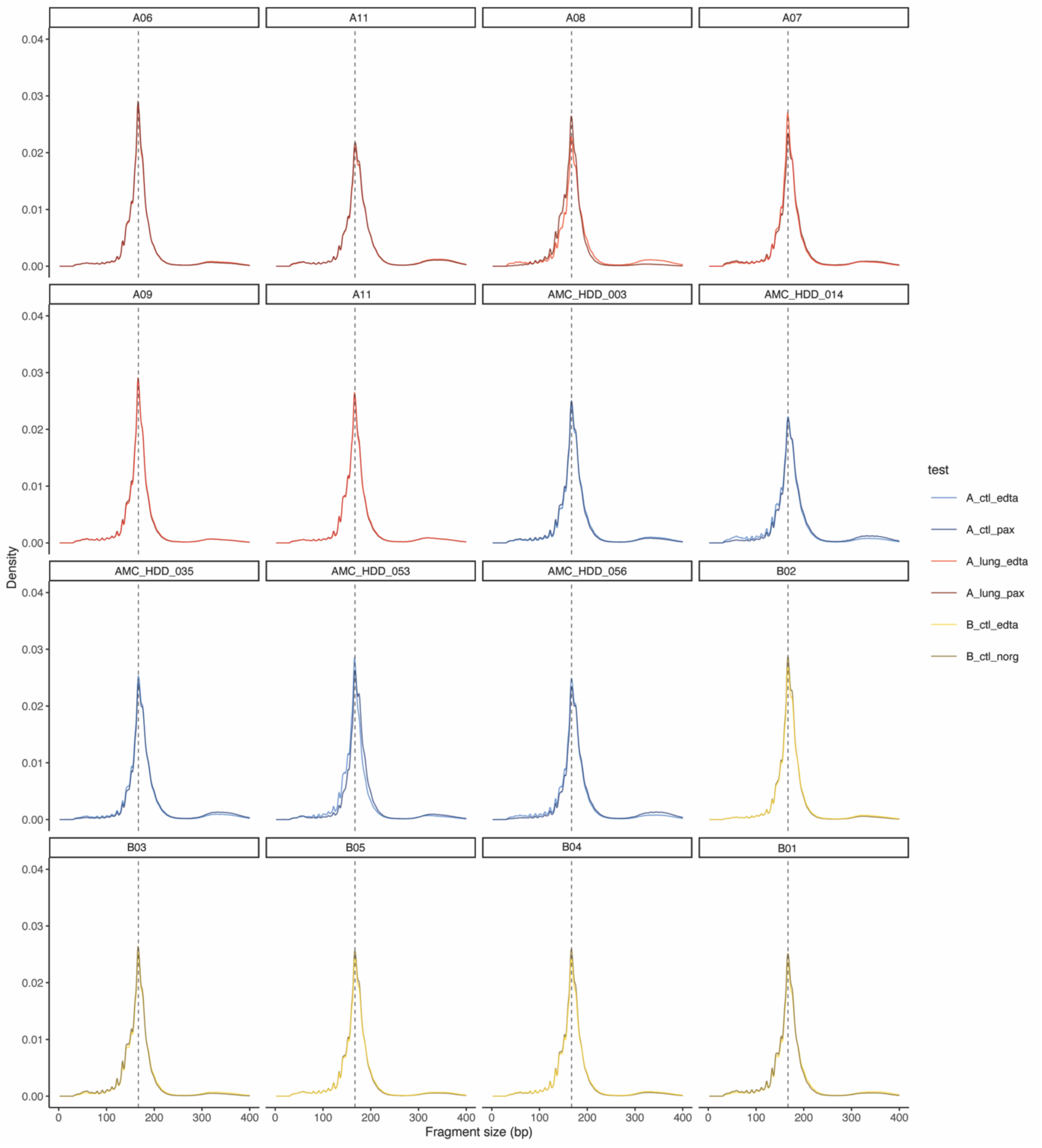


**Suppl. Figure 2: cfDNA fragment size distribution of the plasma samples collected either in EDTA or stabilizing tubes.** Samples are paired by patient. Colors indicate the origin of the samples (A or B), whether the samples are control or lung cancer and the type of tube used. In blue are healthy controls from center A, in red are lung cancer cases from center A and in yellow are healthy controls from center B.

### Suppl. Figure 3

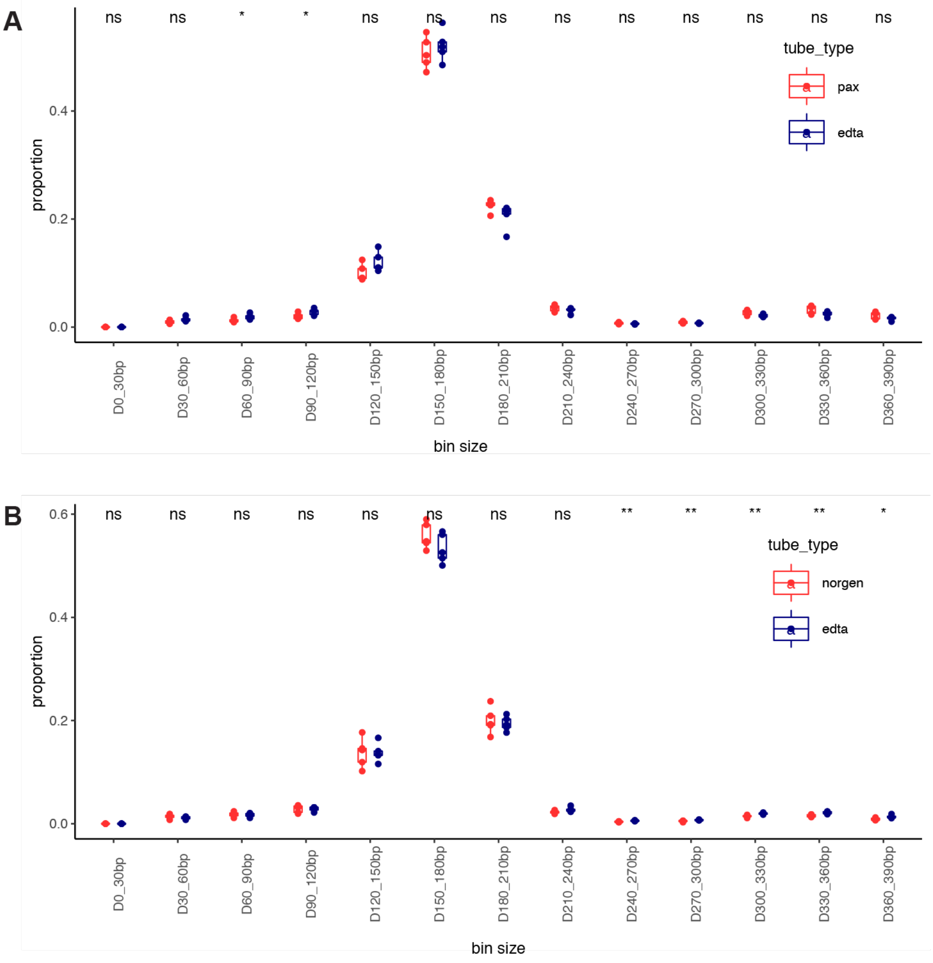


**Suppl. Figure 3: Fragmentation analysis per 30bp size bin. A)** Proportion of fragment sizes grouped per collection tube type in samples collected at clinical center A. **B)** Proportion of fragment sizes grouped per collection tube type in samples collected at clinical center B.

**Suppl. Figure 4**

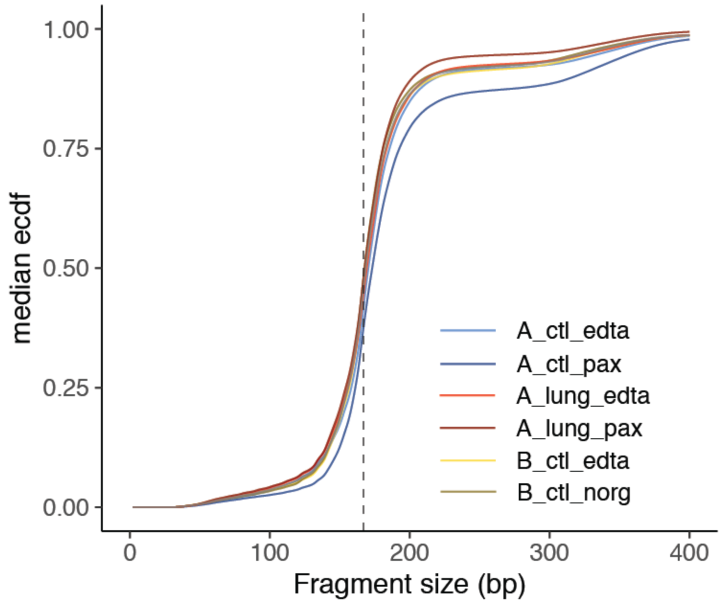


**Suppl. Figure 4 Median empirical cumulative distribution function depending on clinical center (clinical center A and B), sample type (ctl=healthy individual, lung = lung cancer) and collection tube (edta= EDTA, norg = NORGEN, pax = PAXgene).**

### Suppl. Figure 5

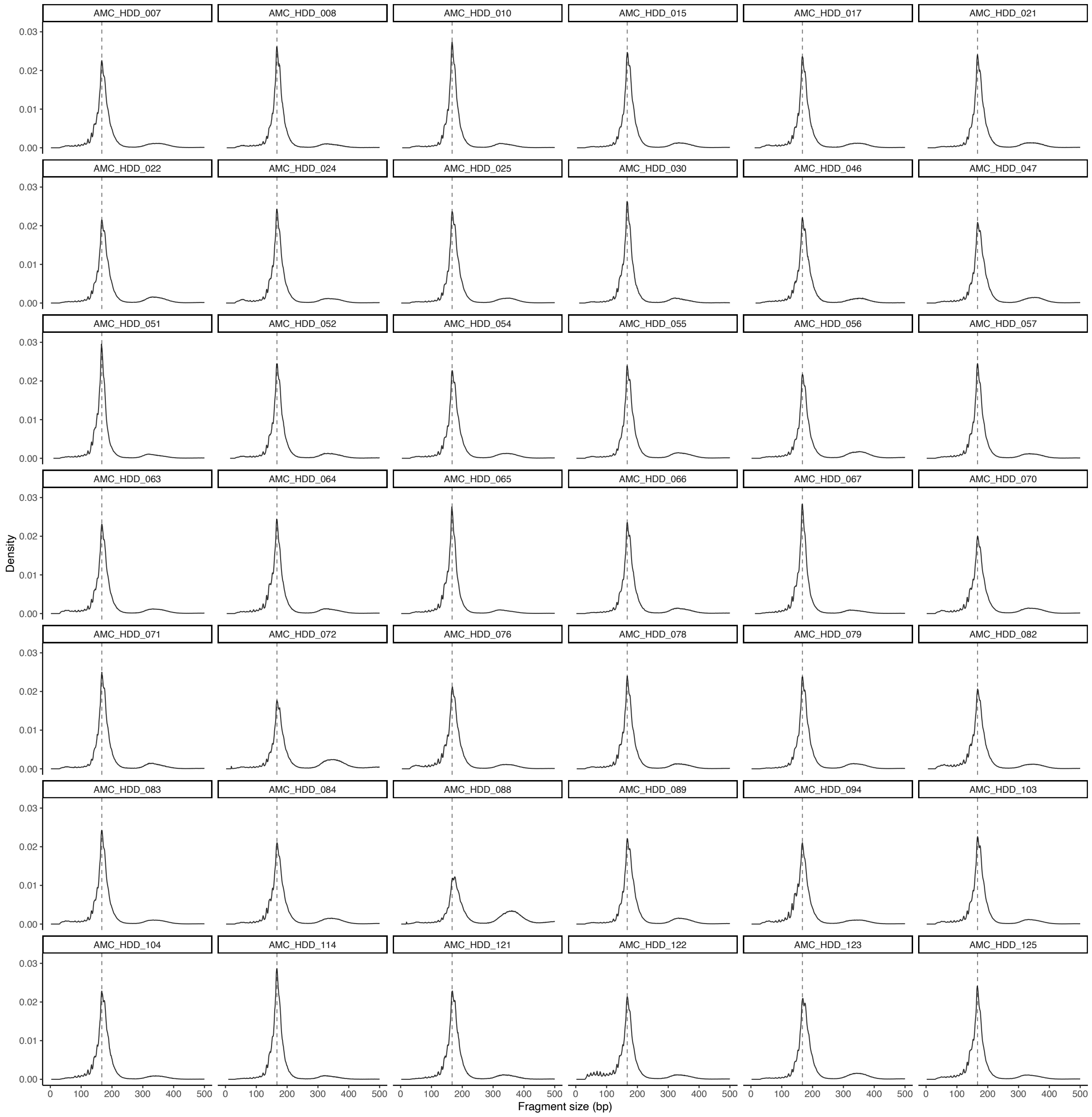


**Suppl. Figure 5: cfDNA fragments size distribution for the samples included in the comparison of physiological variables.** The size distribution is facetted for each sample.

### Suppl. Figure 6

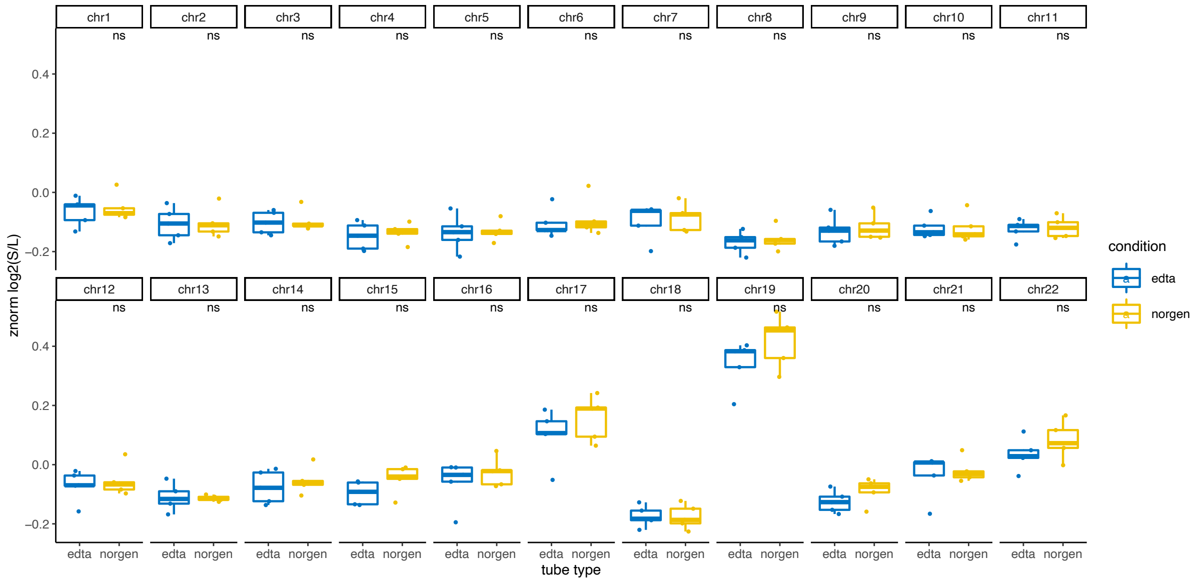


**Suppl. Figure 6: Genome-wide cfDNA fragmentation patterns grouped per collection tube in healthy cases collected at clinical center B.** Paired Wilcoxon rank sum test is applied to test for differences between EDTA – norgen using default statistical significance rank levels.

### Suppl. Figure 7

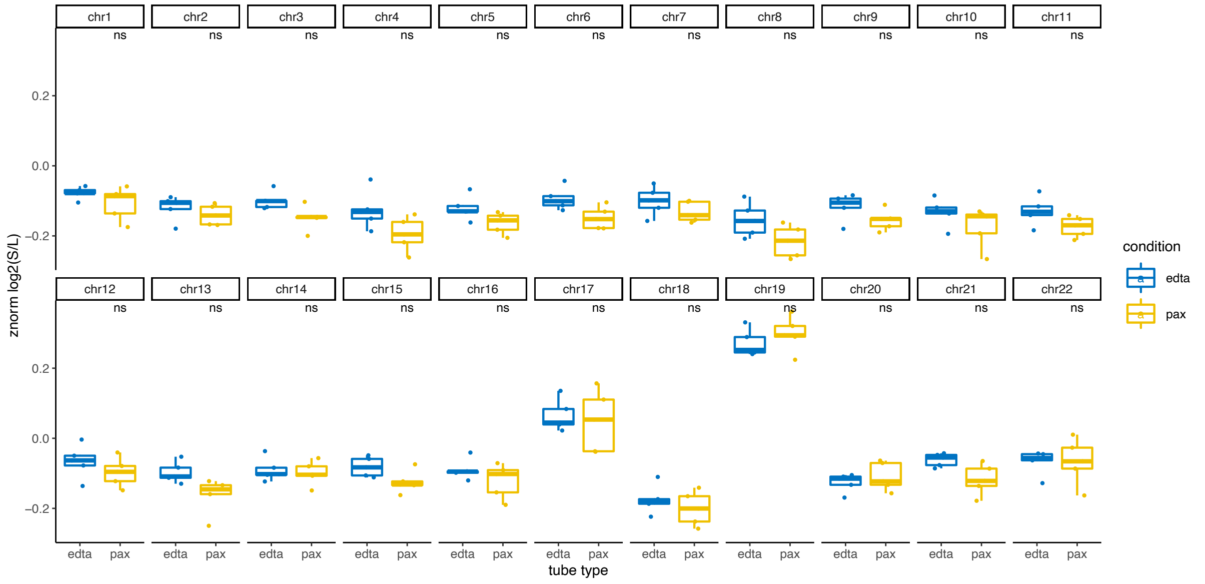


**Suppl. Figure 7: Genome-wide cfDNA fragmentation patterns grouped per collection tube in healthy cases collected at clinical center A.** Paired Wilcoxon rank sum test is applied to test for differences between EDTA – pax using default statistical significance rank levels.

### Suppl. Figure 8

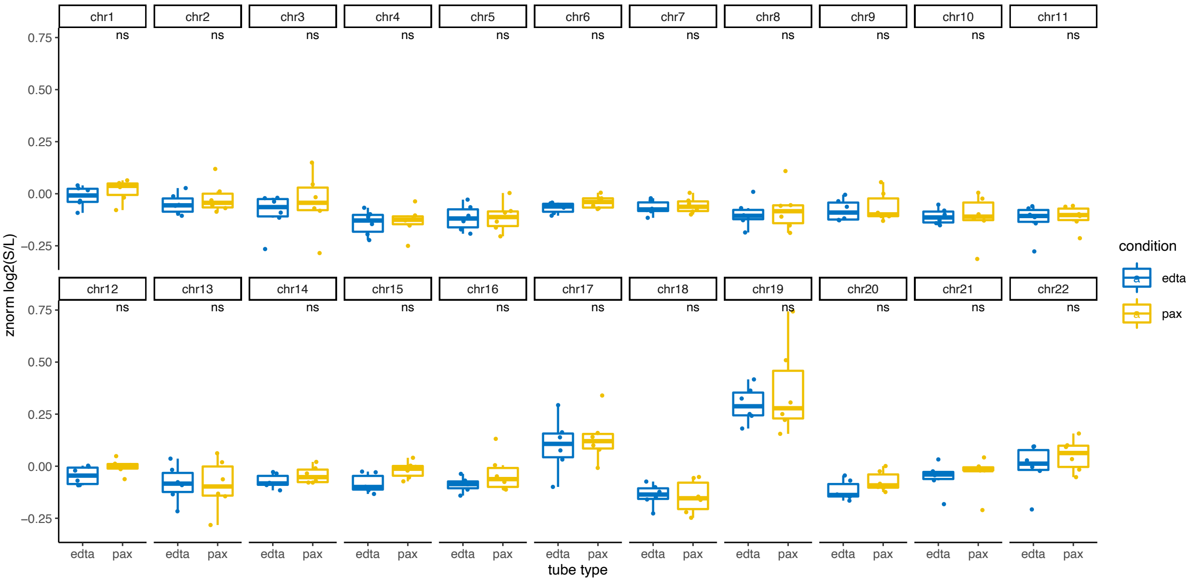


**Suppl. Figure 8: Genome-wide fragmentation patterns grouped per collection tube in lung cancer cases collected at clinical center A.** Paired Wilcoxon rank sum test is applied to test for differences between EDTA – pax using default statistical significance rank levels.

### Suppl. Figure 9

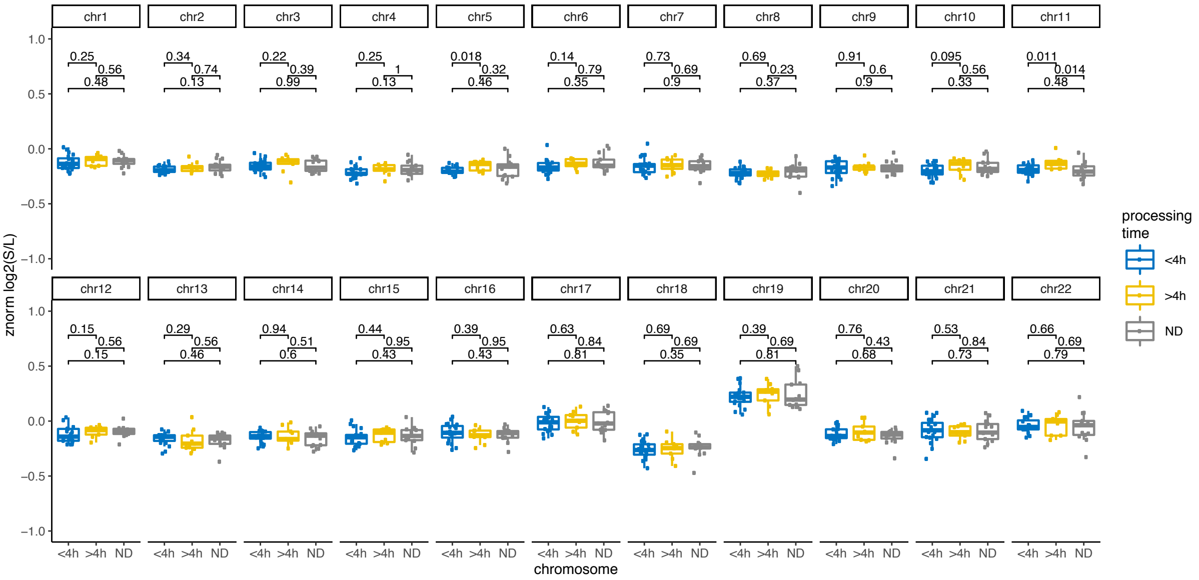


**Suppl. Figure 9: Genome-wide fragmentation patterns grouped per processing time group.** Multivariate comparison is performed using Kruskal-Wallis test, corresponding p values are depicted.

### Suppl. Figure 10

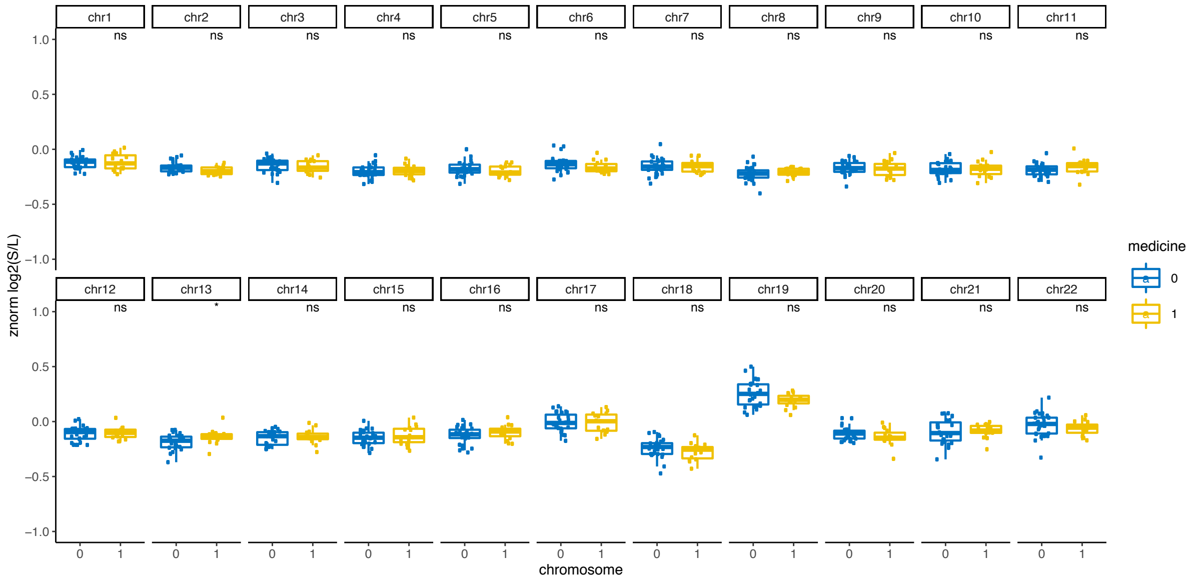


**Suppl. Figure 10: Genome-wide fragmentation patterns grouped for medicine use.** Wilcoxon rank sum test is applied to test for differences between the groups, default statistical significance levels are depicted.

### Suppl. Figure 11

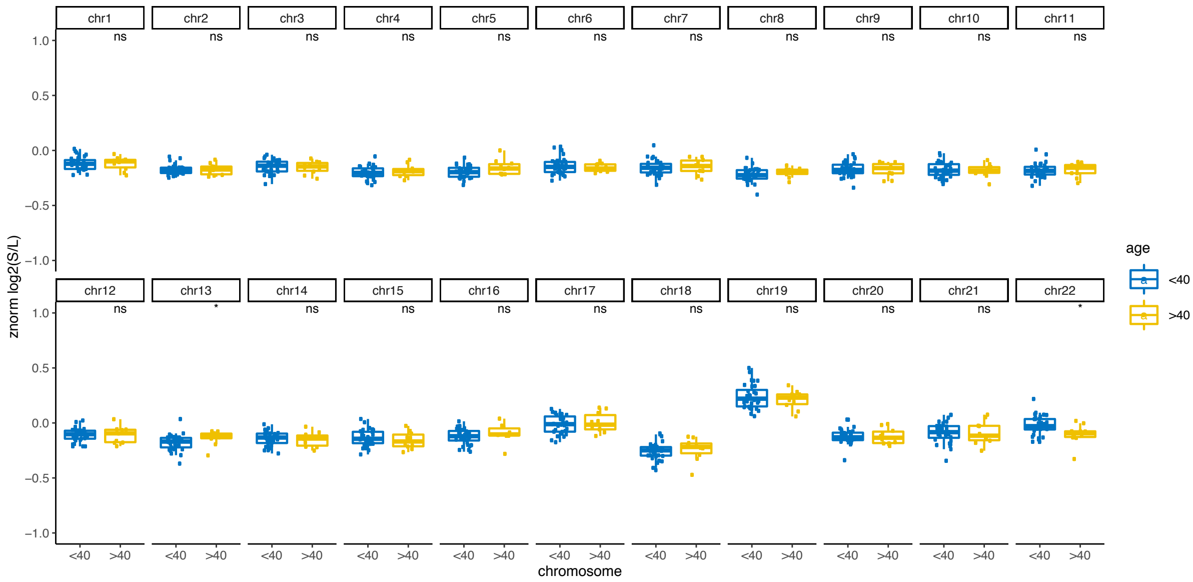


**Suppl. Figure 11: Genome-wide fragmentation patterns grouped by age group.** Wilcoxon rank sum test is applied to test for differences between the groups, default statistical significance levels are depicted.

### Suppl. Figure 12

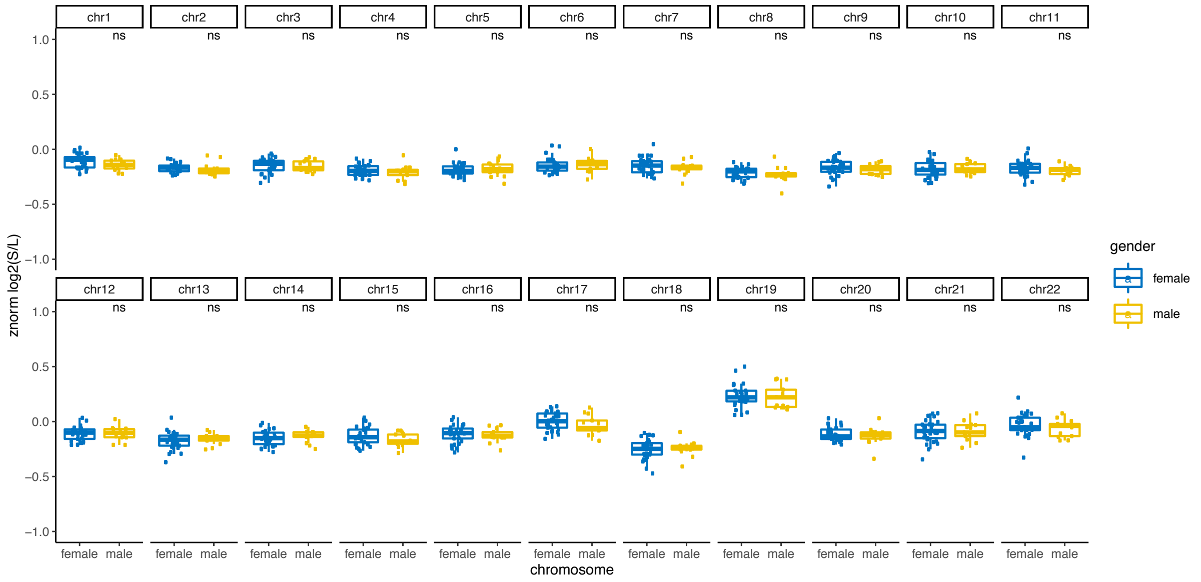


**Suppl. Figure 12: Genome-wide fragmentation patterns grouped by gender.** Wilcoxon rank sum test is applied to test for differences between the groups, default statistical significance levels are depicted.

### Suppl. Figure 13

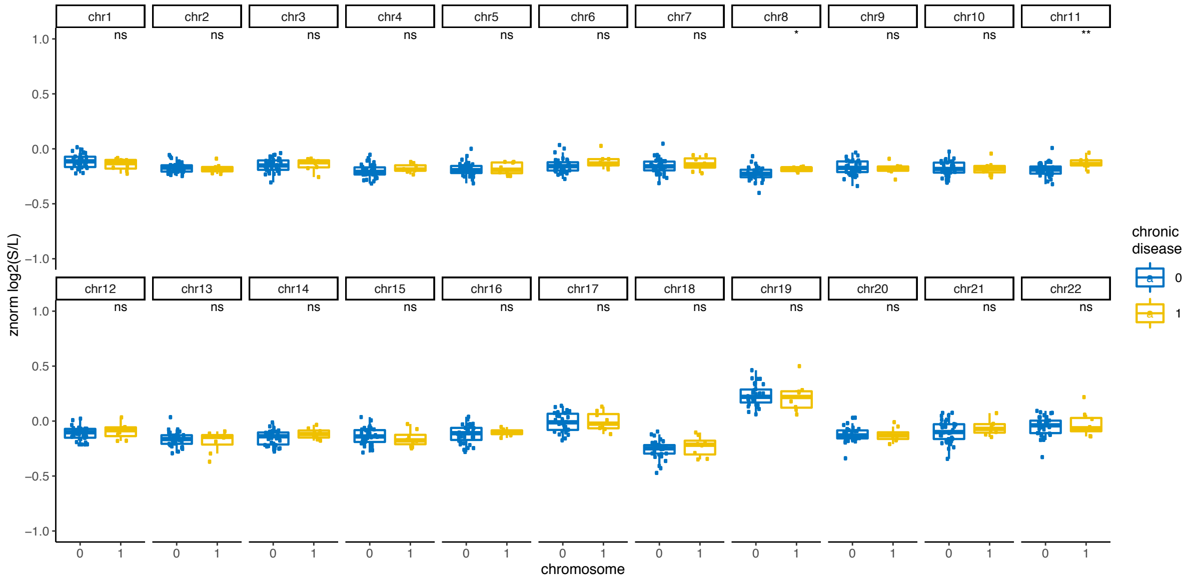


**Suppl. Figure 13: Genome-wide fragmentation patterns grouped by presence of chronic disease.** Wilcoxon rank sum test is applied to test for differences between the groups, default statistical significance levels are depicted.

**Suppl. Figure 14**

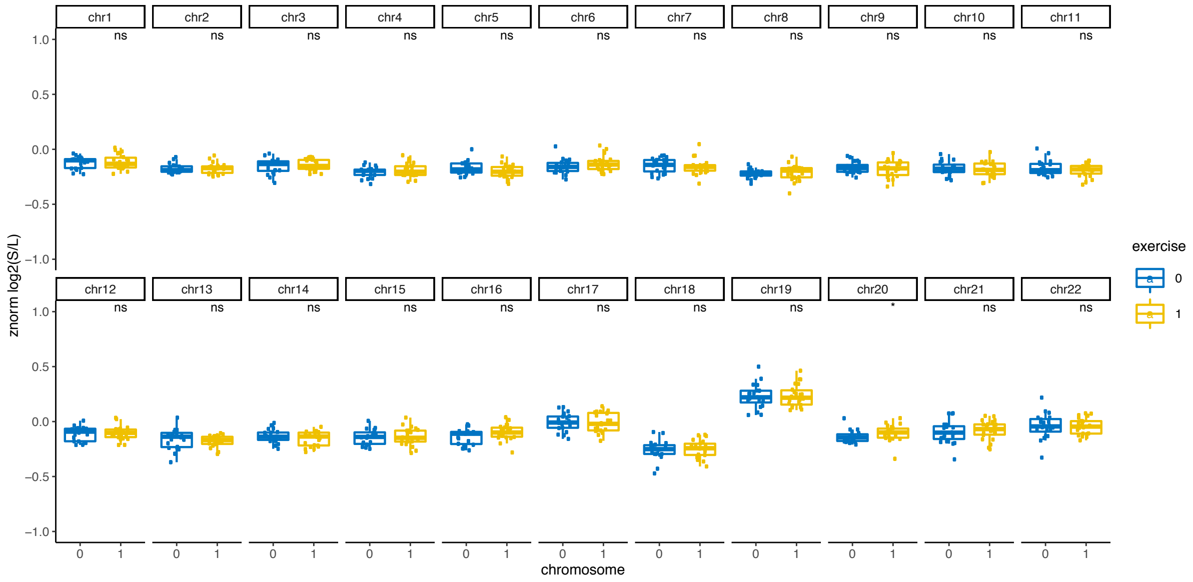


**Suppl. Figure 14: Genome-wide fragmentation patterns grouped by exercise.** Wilcoxon rank sum test is applied to test for differences between the groups, default statistical significance levels are depicted.

**Suppl. Figure 15**

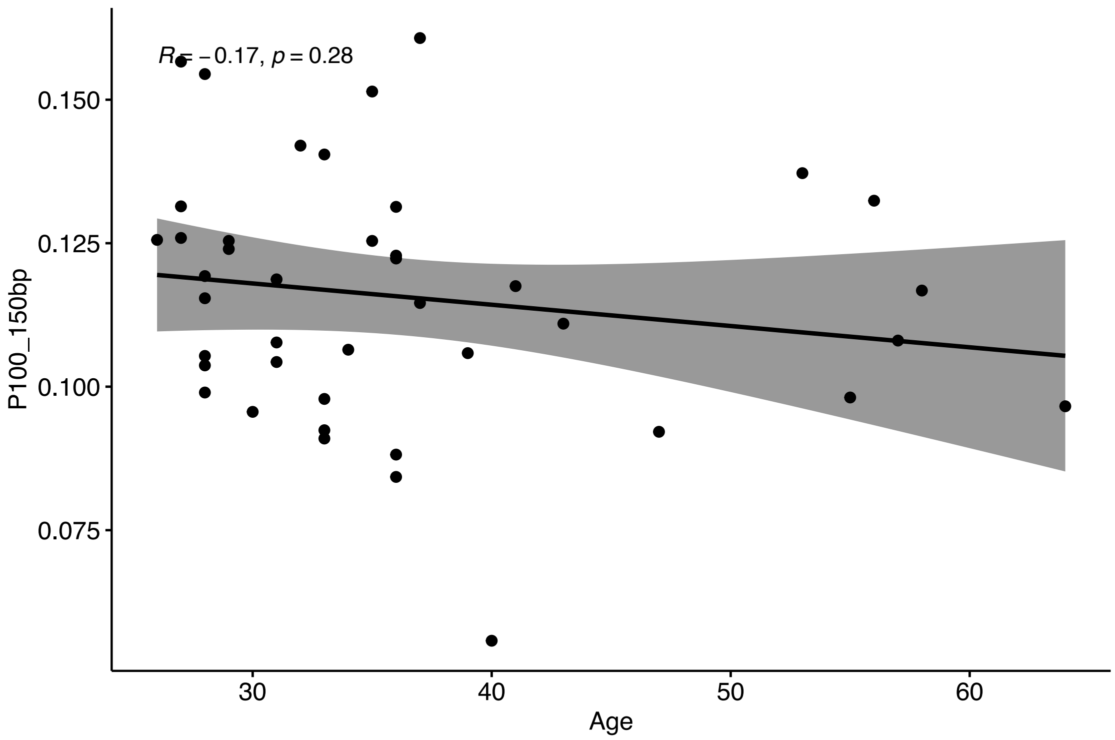


**Suppl. Figure 15: Spearman correlation between Age and P100_150bp, including fitted regression line. Correlation coefficient and p-value are depicted in the upper left corner.**

**Suppl. Figure 16**

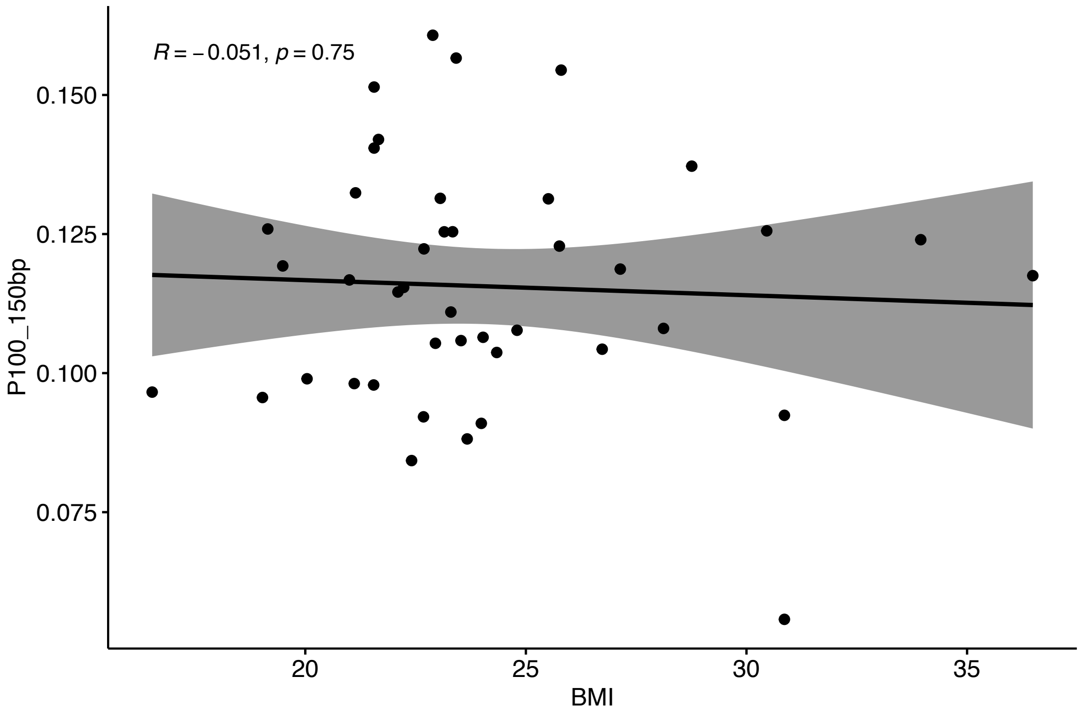


**Suppl. Figure 16: Spearman correlation between BMI and P100_150bp, including fitted regression line. Correlation coefficient and p-value are depicted in the upper left corner.**
